# Differential impact of ageing on reward-driven changes in motor control

**DOI:** 10.64898/2026.08.24.746500

**Authors:** Ahmad Alghamdi, Joseph M. Galea

**Affiliations:** School of Psychology, University of Birmingham, Birmingham, UK; Department of Physical Therapy, College of Applied Medical Sciences, Imam Abdulrahman bin Faisal University, Dammam, Saudi Arabia

**Keywords:** Reward, Ageing, Motor control, Movement vigour, Action selection, Reaching

## Abstract

**Background:** Reward can influence both the selection and execution of goal-directed actions. Healthy ageing is associated with changes in reward processing, raising the possibility that reward effects on motor control may be reduced in older adults.

**Objective:** This study examined how monetary reward affects action execution and action selection during reaching movements and whether these effects differ between younger and older adults.

**Methods:** 28 younger adults and 28 older adults performed a reward-based reaching task. Behaviourally non-distracted trials were used to assess action execution, whereas distractor-containing trials were used to assess action selection. Outcomes included maximum velocity, movement time, endpoint error, reaction time, and selection accuracy.

**Results:** Reward increased maximum velocity and reduced movement time in both age groups without increasing error. These reward-related changes in movement vigour were larger in younger adults. During action selection, reward shortened reaction time but reduced selection accuracy in both groups, indicating faster but less accurate responses. The reward-related changes in reaction time and selection accuracy did not differ significantly between age groups.

**Conclusion:** Ageing did not produce a uniform reduction in reward responsiveness. Instead, ageing attenuated reward-driven movement invigoration, while reward-related changes in action-selection behaviour were similar across age groups. These findings may inform the design of reward-based interventions that promote movement vigour without encouraging speed at the expense of accurate action selection.

## Introduction

Reward can strongly influence motor behaviour. When rewards are available, people often respond quicker, move faster, and perform more accurately. These effects have been demonstrated across different motor tasks, including reaching, saccades, and sequence learning (Klein et al. 2012; Manohar et al. 2015; Codol et al. 2020; Sporn et al. 2022). Reward may therefore improve motor performance by influencing both the selection of an appropriate action and the execution of the selected movement.

Action selection and action execution are related but distinct components of motor control. Action selection refers to the process of choosing an appropriate response from competing alternatives, whereas action execution refers to the generation and control of the selected movement. Previous work suggests that reward can influence both components. In action-selection tasks, reward can reduce reaction time and selection errors, particularly when participants must choose between competing response options (Klein et al. 2012; Manohar et al. 2015; Codol et al. 2020). In action-execution tasks, reward can increase movement vigour, for example by increasing maximum velocity or reducing movement time, whilst maintaining or even improving accuracy (Takikawa et al. 2002; Codol et al. 2020; Galaro et al. 2019). However, most studies have examined these effects in younger adults.

Understanding how reward influences motor control in older adults is important for both theory and rehabilitation. Healthy ageing is associated with changes in motor control, including slower reaction times, longer movement times, and changes in movement strategy (Yan et al. 1998; Seidler et al. 2010; Woods et al. 2015). Ageing is also associated with changes in reward processing, although these changes may differ across behavioural domains. In decision-making and action-selection tasks, older adults often show reduced sensitivity to potential rewards. For example, ageing has been associated with altered reward prediction-error signalling, and dopaminergic modulation can restore aspects of reward learning in some older adults (Chowdhury et al. 2013). Large-scale behavioural studies also suggest that ageing reduces the tendency to approach or select actions associated with potential reward, both in economic decision-making and motor decision-making tasks (Rutledge et al. 2016; Chen et al. 2018). However, evidence from motor-vigour tasks is more mixed. Recent studies suggest that older adults can still use reward expectations to increase response vigour or movement tempo, particularly when task demands are explicit or adapted to individual performance (Hird et al. 2022; Tecilla et al. 2023). Together, these findings suggest that healthy ageing may not produce a uniform reduction in reward sensitivity. Instead, ageing may affect some reward-guided aspects of motor behaviour more than others.

The present study examined the effect of monetary reward on action selection and action execution in younger and older adults using a reaching paradigm previously shown to be sensitive to reward-based changes in motor performance (Codol et al. 2020). Distractor-containing trials were used to assess action selection because participants had to suppress a response to an initial distractor and select the correct target. Behaviourally non-distracted trials, including distractor-free trials and correctly selected distractor-containing trials, were used to assess action execution. This design allowed us to test whether ageing reduces reward-related effects across both components of motor control or affects action selection and action execution differently.

We tested two main hypotheses. First, based on previous work showing beneficial effects of reward on motor performance, we predicted that reward would improve both action selection and action execution. Specifically, we expected reward to improve the speed and accuracy of both distractor-containing (faster reaction times, reduced selection errors) and distractor-free (faster movement time, improved accuracy) trials. Second, based on evidence that ageing is associated with reduced sensitivity to reward, we predicted that reward-related benefits would be smaller in older adults than in younger adults.

## Methods

### Participants

Twenty-eight younger adults (aged 18–25 years, mean age 20 years, 7 men and 21 women) and 28 older adults (aged 40–81 years, mean age 59 years, 18 men and 10 women) took part in the study. Older adults were recruited from a volunteer pool at the University of Birmingham, while younger adults were undergraduate students at the University of Birmingham. Participants were compensated £7.50 per hour plus an additional reward based on their performance. All participants were free from medical or psychiatric conditions that could affect their ability to perform the motor task. The study was approved by the University of Birmingham STEM Ethics Committee (ERN_09-528AP36). All participants provided written informed consent before participation, and the study was conducted in accordance with the Declaration of Helsinki. No formal a priori power analysis was performed. Instead, the sample size was based on previous behavioural work from the laboratory using the same reward-based reaching paradigm, particularly Codol et al. (2020), and on the feasibility of recruiting healthy older adults.

### Task design

The study used the reward-based reaching task described by Codol et al. (2020), implemented with a KINARM end-point robotic device (BKIN Technologies, Ontario, Canada). During the task, participants held a robotic handle that moved horizontally in front of them. Direct vision of the hand was occluded by a horizontal mirror that reflected a screen positioned above it. A white cursor displayed on the screen represented hand position. The screen had a refresh rate of 60 Hz, and kinematic data were recorded at 1 kHz. At the start of each trial, the robotic handle positioned the participant’s hand 4 cm from a fixed home position. A 2-cm-diameter home position then appeared on the screen. Its colour indicated the reward condition: blue or green for no-reward and reward trials, respectively. The potential reward was displayed below the home position as either 0p or 50p.

The task included distractor-free and distractor-containing trials. Distractor-containing trials probed action selection, whereas distractor-free trials allowed direct execution of the reaching movement. To begin each trial, participants moved the cursor to the centre of the home position. A 2-cm-diameter target then appeared 10 cm from the home position. In distractor-free trials, the target had the same colour as the home position, and participants were instructed to reach toward it as quickly as possible and stop on it. In distractor-containing trials, an initial target of a different colour appeared and served as a distractor. Participants were instructed to ignore it and wait for the correct target, which matched the colour of the home position. Reaching toward the distractor resulted in no monetary reward for that trial. Targets appeared at one of four locations positioned at 45° intervals around the midline of the workspace, spanning 135° (Fig. 1a). Luminance-adjusted colours were used to minimise differences in visual detectability (Codol et al. 2020).

**Fig 1.**
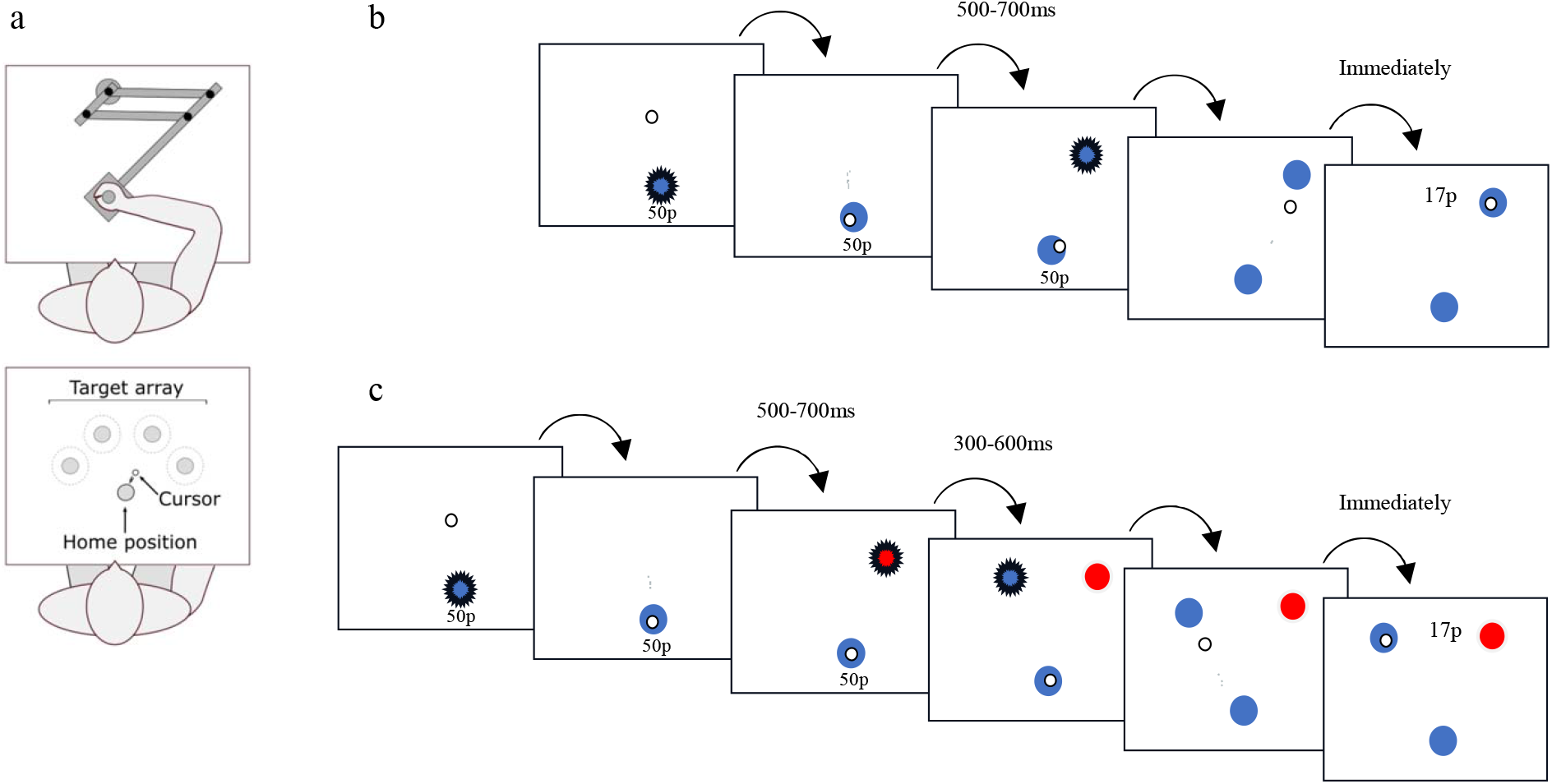
Reaching paradigm. (a) Participants used a robotic manipulandum to reach one of four targets. (b) Distractor-free trial. Participants reached toward a single target and were rewarded according to reaction time plus movement time. (c) Distractor-containing trial. A distractor appeared before the correct target, and participants were instructed to wait for the correct target before reaching.

Participants were informed that monetary reward depended on reaction time and movement time and accumulated throughout the experiment. They were also told that endpoint position was not important provided that the movement ended within 4 cm of the target centre.

The onset of the first target, whether a correct target or a distractor, was sampled from a uniform distribution 500–700 ms after the participant entered the home position. In distractor-containing trials, the correct target appeared 300–600 ms after distractor onset. Once movement velocity fell below 0.03 m/s, the endpoint was recorded and the monetary gain was displayed at the centre of the workspace. After 500 ms, the robotic arm returned the participant’s hand to a position 4 cm from the home position. Participants first completed 12 baseline trials without distractors or reward, followed by 240 experimental trials divided into 24 blocks of 10 trials. Reward and no-reward blocks alternated. Each block contained four distractor-containing trials and six distractor-free trials presented in random order. The starting block condition was counterbalanced across participants.

A closed-loop reward function adapted the reward thresholds to each participant’s recent performance. This individualised the level of challenge and reduced the influence of baseline differences in reaction time and movement speed across participants (Berret et al. 2018; Reppert et al. 2018). The reward function was defined as follows:

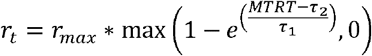

where r_max_ represented the highest possible reward value that could be obtained during a trial. The *MTRT* was the sum of the “total reaction time” and the “movement time”, and τ1 and τ2 were adjustable parameters that depended on the participant’s performance. Specifically, τ1 was calculated as the average of the 3rd and 4th fastest *MTRT*s from the last 20 trials, while τ2 was calculated as the median of the 16th and 17th fastest *MTRT*s from the last 20 trials. At the beginning of each training block, τ1 was set to 400 ms and τ2 to 800 ms, and τ1 was always less than τ2 and both were less than 900 ms. All reward values were rounded up to the nearest penny to ensure that only whole penny values were displayed.

### Data analysis

Distractor-containing trials were manually classified as behaviourally distracted or non-distracted (Fig. 2). Distractor-free trials were automatically classified as non-distracted. A distractor-containing trial was classified as non-distracted when the participant initiated the reach toward the correct target. It was classified as distracted when the participant moved toward the distractor or initially moved toward the distractor and then corrected the movement toward the correct target.

**Fig 2.**
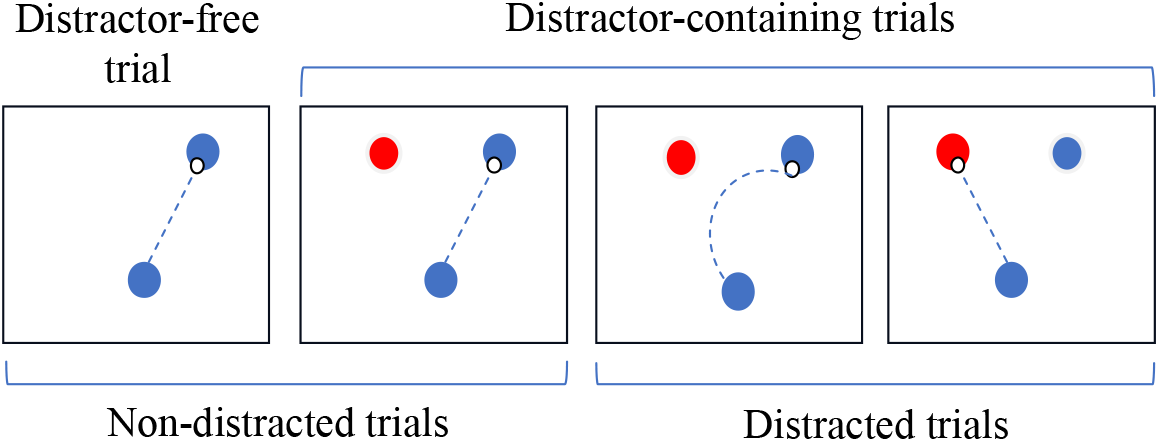
Schematic of trial classification. Distractor-free trials were classified as non-distracted. Distractor-containing trials were classified as non-distracted when the reach was initiated toward the correct target and as distracted when the reach was directed toward the distractor or was corrected after initial movement toward it.

Action selection was assessed using selection accuracy and reaction time. Selection accuracy was defined as the percentage of distractor-containing trials in which the participant initiated the reach toward the correct target. For correctly selected distractor-containing trials, reaction time was defined as the interval from correct-target onset to the point at which the hand moved more than 2 cm from the centre of the home position. For behaviourally distracted trials, reaction time was defined as the interval from distractor onset to the point at which the hand moved more than 2 cm from the centre of the home position.

Action execution was assessed in all behaviourally non-distracted trials, including distractor-free trials and correctly selected distractor-containing trials. The execution outcomes were maximum velocity, movement time, and radial endpoint error. Maximum velocity was the highest tangential hand velocity during the reach. Movement time was the interval from movement onset, defined as the point at which the hand moved more than 2 cm from the centre of the home position, until movement velocity fell below 0.03 m/s. Radial endpoint error was the distance between the centre of the target and the movement endpoint.

Trials were excluded from the kinematic and reaction-time analyses if reaction time was below 200 ms or above 1000 ms, or if a behaviourally non-distracted trial had a radial endpoint error greater than 3 cm. Selection accuracy was calculated from all distractor-containing trials. Overall, 173 of 13,440 experimental trials (1.29%) met at least one exclusion criterion.

### Statistical analysis

Separate 2 × 2 mixed-design analyses of variance (ANOVAs) were conducted for maximum velocity, movement time, radial endpoint error, reaction time, and selection accuracy. Reward condition (reward vs no reward) was the within-subject factor, and Age group (younger vs older) was the between-subject factor. Partial eta squared (ηp^2^) is reported as the effect-size measure.

Where a significant Reward × Age group interaction was detected, follow-up paired-samples t-tests compared the reward and no-reward conditions within each age group, and independent-samples t-tests compared the age groups separately within each reward condition. As significant interactions were observed for two outcomes, four condition-specific age-group comparisons were performed and evaluated using a Bonferroni-adjusted significance threshold of p < 0.0125.

To quantify each significant interaction directly, a reward-effect score was calculated for each participant as mean performance in the reward condition minus mean performance in the no-reward condition. Reward-effect scores were compared between age groups using independent-samples t-tests. These comparisons were evaluated at α = 0.05, and 95% confidence intervals are reported. All tests were two-sided and conducted in MATLAB R2026a (MathWorks, Natick, MA, USA).

## Results

### Monetary Reward Invigorated Action Execution in Both Age Groups

The mixed-design ANOVA showed that monetary reward invigorated action execution in both older and younger adults, as reflected by increased maximum velocity and reduced movement time (Fig. 3; Table 1). There was a significant main effect of Reward on maximum velocity, F(1, 54) = 91.20, p < 0.001, ηp^2^ = 0.628, with no main effect of Age group, F(1, 54) = 0.91, p = 0.343, ηp^2^ = 0.017. The Reward × Age group interaction was significant, F(1, 54) = 6.61, p = 0.013, ηp^2^ = 0.109. Paired-samples t-tests confirmed significant reward-related increases in maximum velocity in both older adults, t(27) = 5.53, p < 0.001, and younger adults, t(27) = 7.81, p < 0.001 (Fig. 3a; Table 1). Movement time showed a similar pattern, with a main effect of Reward, F(1, 54) = 65.61, p < 0.001, ηp^2^ = 0.549, no main effect of Age group, F(1, 54) = 1.37, p = 0.246, ηp^2^ = 0.025, and a significant Reward × Age group interaction, F(1, 54) = 8.54, p = 0.005, ηp^2^ = 0.137. Paired-samples t-tests confirmed significant reward-related reductions in movement time in older adults, t(27) = ™4.24, p < 0.001, and younger adults, t(27) = −6.96, p < 0.001 (Fig. 3b; Table 1). As the Reward × Age group interactions were significant for maximum velocity and movement time, we next examined the reward-related changes within each age group and compared their magnitude between younger and older adults.

**Table 1.** Descriptive statistics and mixed-design ANOVA results. Values are mean (SD). p-values are shown for the main effects of Reward and Age group and for the Reward × Age group interaction. The units cm/s, ms, and mm represent centimetres per second, milliseconds, and millimetres, respectively

|  | Mean (SD) |  |  |  | p (Main effects) |  | p (Interaction) |
| --- | --- | --- | --- | --- | --- | --- | --- |
|  | Older |  | Younger |  | Reward | Age |  |
| Action execution | Reward | No reward | Reward | No reward |  |  |  |
| Maximum velocity (cm/s) | 92.0 (17.4) | 79.1 (17.0) | 92.8 (16.8) | 70.5 (15.3) | < 0.001 | 0.343 | 0.013 |
| Movement time (ms) | 404 (61) | 448 (81) | 397 (38) | 492 (77) | < 0.001 | 0.246 | 0.005 |
| Radial endpoint error (mm) | 11.04 (1.29) | 10.89 (1.15) | 11.64 (1.39) | 11.74 (1.38) | 0.861 | 0.028 | 0.352 |
| Action selection |  |  |  |  |  |  |  |
| Reaction time (ms) | 451 (68) | 490 (71) | 394 (40) | 447 (57) | < 0.001 | 0.002 | 0.180 |
| Selection accuracy (%) | 69.7 (25.3) | 89.8 (9.0) | 70.6 (20.4) | 92.3 (7.5) | < 0.001 | 0.662 | 0.754 |

**Fig 3.**
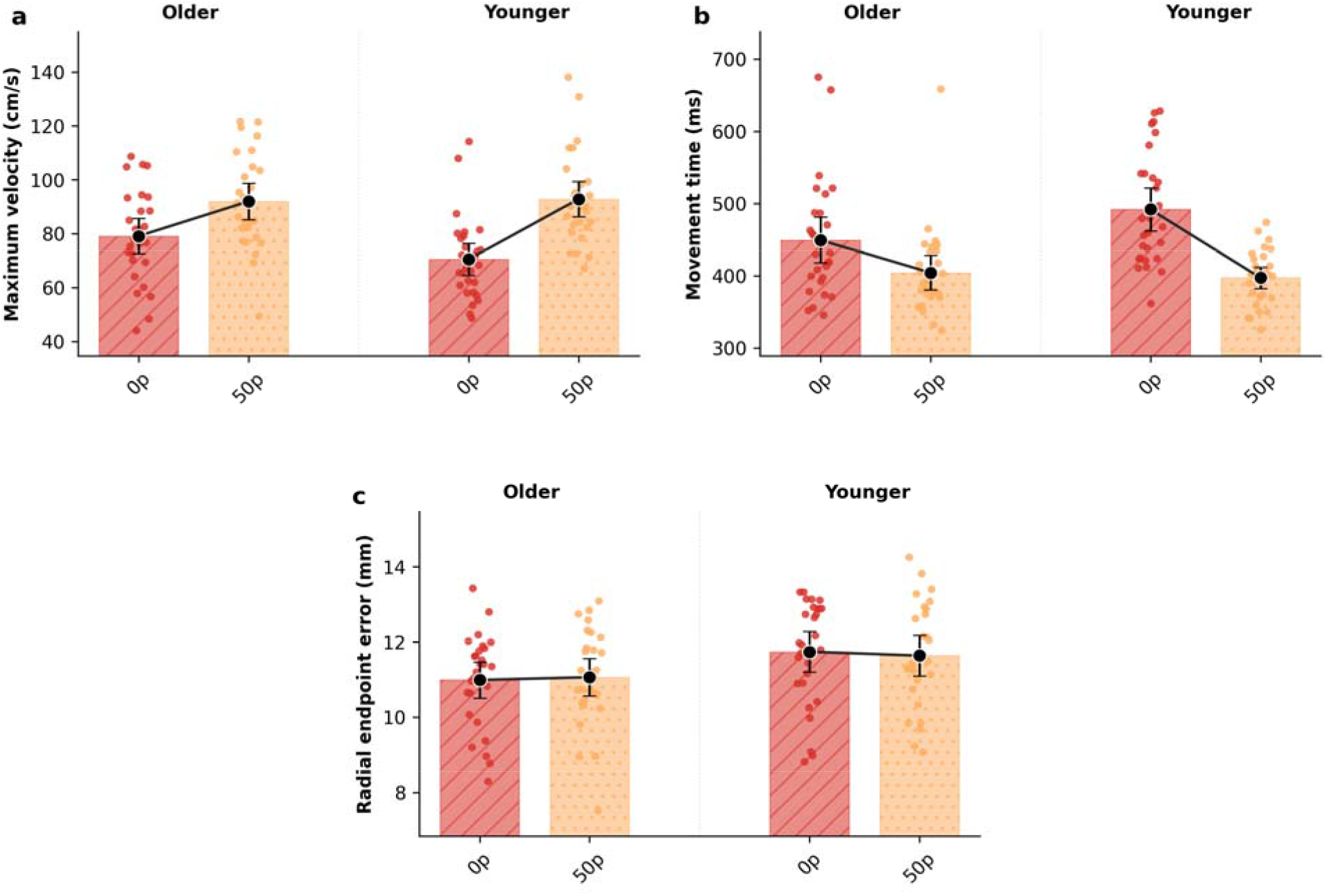
Effects of monetary reward on action execution in older and younger adults. (a) Maximum velocity, (b) movement time, and (c) radial endpoint error. Bars represent group means in the no-reward (0p) and reward (50p) conditions, coloured dots represent individual participants, black circles indicate condition means, connecting lines illustrate the within-group change between conditions, and error bars represent 95% confidence intervals

### Reward-Related Action-Execution Changes Were Larger in Younger Adults

Independent-samples t-tests showed no significant differences between age groups using the Bonferroni-adjusted significance threshold of p < 0.0125 for maximum velocity in either the no-reward condition, t(54) = 1.99, p = 0.051, or the reward condition, t(54) = −0.19, p = 0.850. Similarly, no significant age-group differences were found for movement time in either the no-reward condition, t(54) = −2.05, p = 0.046, or the reward condition, t(54) = 0.50, p = 0.622.

However, the significant Reward × Age group interactions indicated that the magnitude of reward-related change differed between younger and older adults. To quantify these interactions directly, reward-effect scores were calculated by subtracting each participant’s mean performance in the no-reward condition from their mean performance in the reward condition (reward minus no reward; Fig. 4). These scores were then compared between age groups using independent-samples t-tests. For maximum velocity, the mean reward effect was 12.9 cm/s in older adults and 22.3 cm/s in younger adults; the younger-minus-older difference was 9.48 cm/s, t(54) = 2.57, p = 0.013, 95% CI [2.09, 16.87]. For movement time, the mean reward effect was −44.4 ms in older adults and −94.6 ms in younger adults; the younger-minus-older difference was −50.2 ms, t(54) = −2.92, p = 0.005, 95% CI [−84.57, −15.76]. Negative movement-time scores indicate that movements were shorter in the reward condition. Thus, reward invigorated action execution in both groups, but the reward-related increase in maximum velocity and reduction in movement time were significantly larger in younger adults.

**Fig 4.**
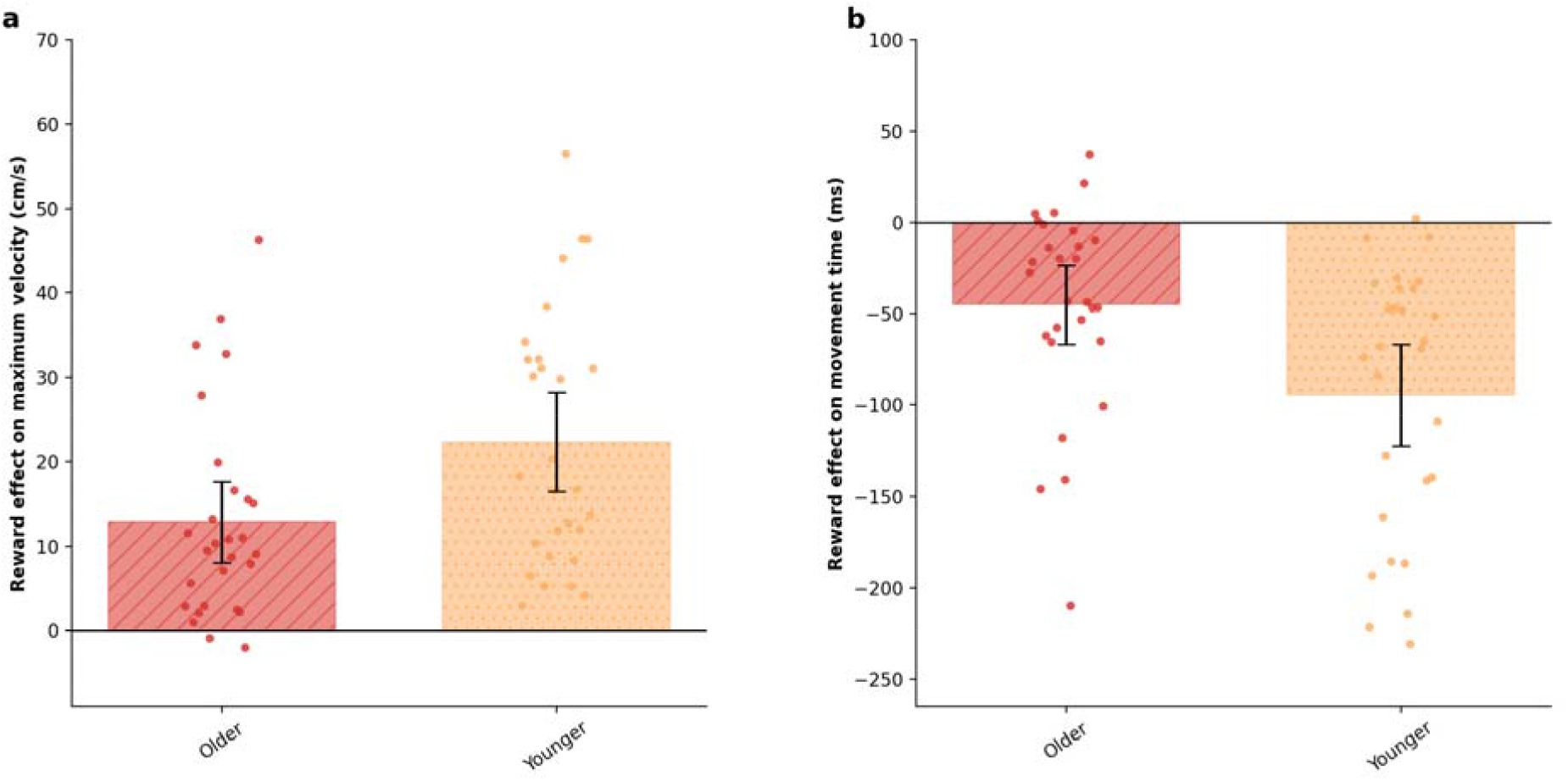
Reward effects on action execution by age group. Reward-effect scores were calculated as reward minus no reward for (a) maximum velocity and (b) movement time. Positive values indicate increased velocity, whereas negative values indicate reduced movement time. Bars show group means, dots show individual participants, and error bars show 95% confidence intervals

### Reward-Driven Invigoration Did Not Compromise Endpoint Accuracy

Radial endpoint error was analysed to determine whether reward-related increases in movement vigour were accompanied by reduced spatial accuracy. Older adults showed lower radial endpoint error than younger adults overall, as indicated by a significant main effect of Age group, F(1, 54) = 5.07, p = 0.028, ηp^2^ = 0.086. However, there was no significant main effect of Reward, F(1, 54) = 0.03, p = 0.861, ηp^2^ < 0.001, and no Reward × Age group interaction, F(1, 54) = 0.88, p = 0.352, ηp^2^ = 0.016. Therefore, despite an overall age-group difference in endpoint accuracy, reward-related increases in movement vigour did not come at the cost of reduced spatial accuracy in either group (Fig. 3c; Table 1).

### Reward Shortened Reaction Time but Reduced Selection Accuracy

Reward shortened reaction time, as shown by a significant main effect of Reward, F(1, 54) = 82.19, p < 0.001, ηp^2^ = 0.604. There was also a significant main effect of Age group, F(1, 54) = 10.66, p = 0.002, ηp^2^ = 0.165, with older adults responding more slowly overall. The Reward × Age group interaction was not significant, F(1, 54) = 1.85, p = 0.180, ηp^2^ = 0.033, indicating that the reward-related reduction in reaction time did not differ significantly between age groups (Fig. 5a; Table 1). In contrast to its effect on reaction time, reward reduced selection accuracy, as indicated by a significant main effect of Reward, F(1, 54) = 67.15, p < 0.001, ηp^2^ = 0.554. Neither the main effect of Age group, F(1, 54) = 0.19, p = 0.662, ηp^2^ = 0.004, nor the Reward × Age group interaction, F(1, 54) = 0.10, p = 0.754, ηp^2^ = 0.002, was significant (Fig. 5b; Table 1). Together, the shorter reaction times and lower selection accuracy indicate that reward shifted both age groups toward faster but less accurate action selection.

**Fig 5.**
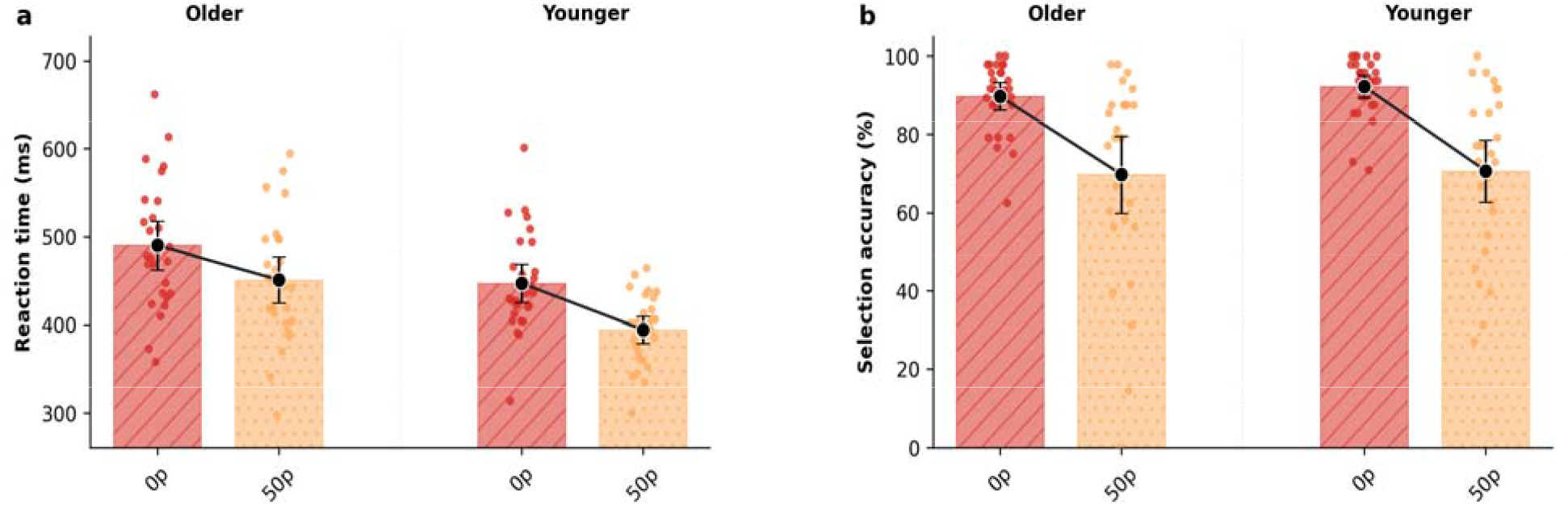
Effects of monetary reward on action selection in older and younger adults. (a) Reaction time and (b) selection accuracy. Bars represent group means in the no-reward (0p) and reward (50p) conditions, coloured dots represent individual participants, black circles indicate condition means, connecting lines illustrate the within-group change between conditions, and error bars represent 95% confidence intervals

## Discussion

The present study examined how monetary reward influenced action selection and action execution in younger and older adults. We predicted that reward would improve speed and accuracy performance across both components of motor control and that ageing would produce a global reduction in these reward-related benefits. The findings only partly supported these predictions. Reward influenced action execution and action selection in dissociable ways. During action execution, reward increased movement vigour, reflected by higher maximum velocity and shorter movement time, without affecting accuracy (endpoint error); however, this reward-driven invigoration was significantly smaller in older adults than in younger adults. During action selection, reward shortened reaction time but unexpectedly reduced selection accuracy, shifting behaviour towards faster but less accurate responses. This reward-induced speed–accuracy trade-off did not differ significantly between age groups. Thus, ageing did not produce the predicted global reduction in reward responsiveness. Instead, it selectively attenuated reward-driven movement invigoration while leaving reward-related adjustments in action-selection behaviour relatively preserved.

### Reward-driven movement invigoration is attenuated in older adults

Reward improved the execution of reaching movements in both younger and older adults. Participants moved faster in rewarded trials than in non-rewarded trials, as shown by increased maximum velocity and reduced movement time. This finding is consistent with previous studies showing that reward can invigorate movement execution in reaching and other motor tasks (Codol et al. 2020; Galaro et al. 2019; Manohar et al. 2015; Sporn et al. 2022). Importantly, reward did not increase radial error, indicating that the increase in execution speed was not achieved at the cost of reduced endpoint accuracy. Thus, reward enhanced movement vigour while preserving spatial accuracy.

Although both age groups showed reward-related invigoration, the magnitude of this effect was smaller in older adults. Younger adults showed a larger reward-related increase in maximum velocity and a larger reward-related reduction in movement time. This finding supports our prediction that reward-related benefits would be reduced in older adults. It also fits with evidence that healthy ageing is associated with changes in reward processing and dopaminergic function (Chowdhury et al. 2013; Eppinger et al. 2012; Rutledge et al. 2016). However, the present study did not directly measure dopamine function or neural activity, so this interpretation should remain cautious. Behaviourally, the key point is that older adults were still sensitive to reward, but the invigorating effect of reward on movement execution was attenuated.

Older adults were also more accurate than younger adults overall, as reflected by lower radial error. This is consistent with previous work suggesting that older adults may adopt more conservative motor strategies, prioritising accuracy and control over speed (Helsen et al. 2016; Lee et al. 2007; Seidler et al. 2010). Therefore, the smaller reward-related increase in movement vigour in older adults may reflect a combination of reduced reward sensitivity and a greater tendency to preserve movement accuracy. However, because reward did not increase radial error in either age group, reward-driven execution invigoration did not produce a conventional speed–accuracy cost in terms of endpoint accuracy.

### Reward speeds action selection but reduces selection accuracy

Reward had a different effect on action selection. In distractor-containing trials, reward shortened reaction time in both age groups, indicating that monetary incentives accelerated response initiation. Older adults were slower overall than younger adults, consistent with previous evidence of age-related slowing in response preparation and selection (Hardwick et al. 2022; Woods et al. 2015). However, the reward-related reduction in reaction time did not differ significantly between age groups. Thus, older adults remained able to use reward cues to speed response initiation despite slower overall reaction times.

Contrary to our initial prediction, reward did not improve selection accuracy. Instead, reward reduced selection accuracy in both age groups. Thus, in distractor-containing trials, reward shifted behaviour toward faster but less accurate responses. This suggests that reward altered the balance between speed and accuracy during action selection. As monetary gain was linked to response speed, participants may have prioritised faster initiation, increasing the likelihood of responding before the correct target was fully selected. However, this interpretation should be treated cautiously because the present task was not designed to directly measure decision thresholds or response urgency.

This finding contrasts with previous work showing that reward can improve action selection in the presence of competing distractors, leading to faster and more accurate responses (Codol et al. 2020; Manohar et al. 2015). The reason for this discrepancy is unclear, particularly because the present study used a closely related task and reward structure. At a minimum, these findings suggest that reward effects on action selection may be less consistent than reward effects on action execution. While reward reliably increased movement vigour in both younger and older adults, its effect on action selection may depend on the balance between speeding responses and maintaining selection accuracy. Future work should directly test how different reward structures, such as rewarding speed, accuracy, or both, influence action selection in younger and older adults.

### Ageing and reward sensitivity across motor-control components

The present findings partially support the hypothesis that ageing reduces reward-related benefits. Ageing attenuated reward-related changes in action execution but did not significantly alter the reward-related pattern observed during action selection, where reward shortened reaction time and reduced selection accuracy in both groups. Thus, ageing did not simply reduce all reward effects; its influence depended on the behavioural component being measured.

This finding fits with previous evidence that ageing can alter reward processing, but also suggests that such changes are not uniform across tasks. Chowdhury et al. (2013) showed that older adults had altered reward prediction-error signalling in the nucleus accumbens, driven particularly by abnormal expected-value representation. Importantly, L-DOPA increased learning rate and improved task performance in some older adults, and this behavioural improvement was linked to restoration of a more canonical reward prediction-error signal. This provides evidence that age-related changes in dopaminergic reward processing can influence how older adults use reward information to guide behaviour. Similarly, large-scale behavioural studies suggest that ageing reduces reward-related approach behaviour. Rutledge et al. (2016) found that risk taking for potential reward decreases across the lifespan, and Chen et al. (2018) showed that ageing reduces Pavlovian attraction toward reward during motor decision-making. These findings support the idea that older adults may show reduced sensitivity to reward when reward must guide learning, choice, or action selection.

However, evidence from motor-vigour tasks is more mixed. Hird et al. (2022) found that average reward rate influenced response vigour similarly in younger and older adults, while Tecilla et al. (2023) showed that trial-by-trial expectations about action–reward contingencies increased motor tempo in both younger and older adults. These findings suggest that reward-related invigoration can be preserved under some conditions, particularly when reward information is explicit or when the task is adapted to individual performance. The present findings add to this literature by showing that, within the same reaching task, reward-related changes in action-selection behaviour did not differ significantly between age groups but reduced reward-driven invigoration of movement execution. This dissociation suggests that the effect of ageing on reward sensitivity depends not only on whether reward is present, but also on the specific motor-control process being measured and on how reward is linked to task performance.

### Limitations and future directions

Two main limitations should be considered. First, the study used monetary reward, but its subjective value may not have been equivalent across age groups. Younger adults were undergraduate students, whereas older adults were recruited from a volunteer pool, and the monetary incentive may therefore have been experienced differently by the two groups. Future studies should directly assess subjective reward value, intrinsic motivation, and task engagement to determine whether these factors explain age-related differences in reward responsiveness.

Second, reward was linked to performance speed, which may have encouraged participants to prioritise rapid responding. This is particularly relevant to the action-selection results, in which shorter reaction times were accompanied by lower selection accuracy. Future studies should compare reward conditions that emphasise speed, accuracy, or both to determine whether the reduction in selection accuracy reflects strategic prioritisation of speed or a broader effect of reward on action selection.

### Implications for reward-based rehabilitation

Despite these limitations, the findings may have implications for the design of reward-based rehabilitation. Reward and punishment can influence motor adaptation after stroke (Quattrocchi et al. 2017), and motivation is an important determinant of engagement in older adults (Hess 2014). The present results show that monetary reward can increase movement vigour in healthy older adults, although the magnitude of this effect was smaller than in younger adults.

Future rehabilitation studies should test whether tailoring incentives to patients’ goals, preferences, and functional outcomes leads to more clinically meaningful changes in motor performance.

The action-selection findings also highlight a potential limitation of speed-focused incentives. Reward shortened reaction time but reduced selection accuracy in both age groups. In a rehabilitation context, prioritising rapid responses could be problematic if it encourages patients to initiate movements before the appropriate action has been reliably selected or planned. Although this possibility requires direct testing in clinical populations, rehabilitation protocols that reward speed should therefore consider reinforcing accuracy, movement quality, and functional success alongside movement vigour.

### Conclusion

In conclusion, monetary reward affected action execution and action selection in distinct ways. Reward increased movement vigour without compromising endpoint accuracy, although this effect was smaller in older adults. During action selection, reward shortened reaction time but reduced selection accuracy, and these reward-related changes did not differ significantly between age groups. Thus, contrary to the predicted global reduction in reward responsiveness, older age was associated specifically with attenuated reward-driven movement invigoration, while reward-related changes in action-selection behaviour were relatively preserved. Future rehabilitation studies should examine reward schedules that promote movement vigour while maintaining accurate action selection.

## Statements and Declarations

## Ethics approval

The study was approved by the University of Birmingham STEM Ethics Committee (ERN_09-528AP36) and was conducted in accordance with the Declaration of Helsinki.

## Consent to participate

Informed consent was obtained from all participants.

## Data availability

The de-identified data, labelled distractor-trial figures, and MATLAB analysis code used in this study are publicly available on the Open Science Framework at https://osf.io/5vynd/.

## Author contributions

Ahmad Alghamdi conducted the study, collected the data, performed the analyses, interpreted the findings, and wrote the original manuscript. Joseph M. Galea supervised the research, provided resources, secured funding for participant compensation, and critically reviewed and revised the manuscript. All authors read and approved the final manuscript.

